# Morphological and MALDI-TOF MS identification of freshwater snails, including intermediate hosts of schistosomes in Gabon

**DOI:** 10.1101/2025.07.14.664644

**Authors:** Lady Charlène Kouna, Sandrine Lydie Oyegue-Liabagui, Nick Chenis Atiga, Alain Prince Okouga, Adama Zan Diarra, Steede Seinnat Ontoua, Gabriel Falque, Stéphane Ranque, Jean Bernard Lekana-Douki

## Abstract

A thorough knowledge of intermediate snail hosts and their geographical distribution is essential to understanding the transmission of schistosomiasis and to inform interventions in Gabon. Accurate identification of both snails species and *Schistosoma* species is therefore essential. The aim of this study was to identify the intermediate hosts of *Schistosoma* spp. (snails) using MALDI-TOF MS.

**Methods:** Snails were collected from various sites across Gabon and were morphologically identified to species or genus level. Snail species were randomly selected, dissected and subjected to MALDI-TOF MS and molecular analysis. Parasites were detected by cercarial emission, followed by qPCR analysis and sequencing.

**Results:** A total of 3,722 snails were collected and morphologically identified as *Bulinus truncatus* (n=1845), *Bulinus forskalii* (n=381), *Gyraulus costulatus* (n=680), *Bellamya* sp. (n=67) and *Melania* sp. (n=749). Of the 3,722 snails collected, 21.2% (n = 791) were selected for further analysis, including *Bu. truncatus* (n = 301), *Bu. forskalii* (n = 158), and *Gy. costulatus* (n=147).

A subset of 65 snails was subjected to molecular identification to confirm morphological classification, using *COI*, *ITS*, and *18S* gene. BLAST analysis of the resulting sequences showed 97–100% identity with reference sequences of *Bu. truncatus*, *Bu. forskalii*, and *Gy. costulatus* available in GenBank. Analysis of the MS spectra from the 791 snails revealed that 78.2% (n = 618) were of high quality, with signal intensities exceeding 3,000 arbitrary units (a.u.). The MALDI-TOF MS reference database was subsequently updated with 12 high-quality spectra, comprising four representative spectra for each snail species. A blind test of 606 remaining spectra against the database. A blind test of the remaining 606 spectra, using the updated MALDI-TOF MS reference database, demonstrated reliable identification of 100% (*Bulinus forskalii*, 147/147; *Gyraulus costulatus*, 158/158) and 96% (*Bulinus truncatus*, 289/301) of the specimens. These results were consistent with morphological identification, with log score values (LSVs) ranging from 1.7 to 2.8. Principal component analysis (PCA) showed that *Bulinus* species clustered according to their geographical origin. Overall, 17,66% of the snails analysed were found to be positive for *Schistosoma* sp. Further analysis, including qPCR and sequencing, revealed that the snails were infected with *Schistosoma haematobium*.

**Conclusion:** We report the first data on the identification of snails by MALDI-TOF MS in Gabon and show that *Bu. truncatus* and *Bu. forskalii* are potential intermediate hosts of *S. haematobium*. We also show a heterogeneous spatiotemporal distribution of snails in south-eastern Gabon. The role of *G. costulatus* in the transmission of schistosome remains unclear.

**Author Summary:** Schistosomiasis is a disease caused by trematodes of the genus *Schistosoma*. Freshwater snails of the genera *Biomphalaria*, *Bulinus* and *Oncomelania* release infectious cercariae, which are transmitted to the final host (human or animal). It is the second most widespread parasitic disease after malaria, and among the most important neglected tropical diseases in the world. Snail identification is extremely important for monitoring snail populations and schistosomiasis. Identification has long relied on morphological criteria and molecular biology, both of which have several drawbacks. However, numerous studies have reported the performance of MALDI-TOF MS, a technology that allows species to be identified based on their proteins, as a reliable, rapid and easy-to-use tool in many fields. Recently, it has been described as an effective tool for snail identification and even for tracing the origin of the intermediate hosts of schistosomiasis. The objective of this study was to accurately identify freshwater snails, including the intermediate hosts of schistosomiasis in Gabon.

## INTRODUCTION

Flukes of the genus *Schistosoma* cause schistosomiasis. This is a debilitating disease y that mainly affects poor populations in tropical and subtropical areas who have limited access to potable water and poor hygiene conditions and whose main activities are carried out in freshwater [1]. It is the second most important parasitic disease after malaria [2–4]. According to the World Health Organization (WHO), in 2021, approximately 251.4 million individuals required treatment for schistosomiasis in 52 countries, predominantly in sub-Saharan Africa, where schistosomiasis was predominantly endemic and it was estimated that 91% of cases occurred [1,5,6]. Gastropods of the genera *Bulinus*, *Oncomelania* and *Biomphalaria* are known to be intermediate hosts for the larval development of parasite species of the genus *Schistosoma* sp., in endemic regions [7,8]. The geographical distribution of trematodes responsible for schistosomiasis is determined by the presence of their intermediate hosts [7]. In the sub-Saharan region of Africa*, Biomphalaria pfeifferi* is the only intermediate host of *Schistosoma mansoni*, while snails of the genus *Bulinus* are involved in the transmission of *Schistosoma haematobium* and *Schistosoma guineensis/intercalatum* [9–12]. The genus *Bulinus* comprises four species: *Bulinus africanus*, *Bu. forskalii*, *Bu. reticulatus* and the *Bu. truncatus*/*tropicus* complex [13]. However, the role of each species in disease transmission varies considerably depending on geographical location [14]. For example, in Gabon, *Bu. truncatus* is involved in the transmission of urinary schistosomiasis (35,36), whereas in Senegal, *Bu. globosus, Bu. senegalensis Bu. umbilicatus* and *Bu. forskalii* are known to be intermediate hosts of *S. haematobium* [11,12].

Effective control of schistosomiasis requires the correct identification of the intermediate snail host [15]. Several methods have been described for identifying snails, including macro and microscopic morphological description, genomic and mitochondrial DNA sequence-based analysis, and proteomic profiling techniques using matrix-assisted laser desorption/ionisation time-of-flight mass spectrometry (MALDI-TOF MS) [7,16–22]. However, accurately identifying snail species based on shell morphology is laborious and requires high-quality specimens and malacological experts with extensive training [16,21,23]. The DNA sequence-based approach overcomes these limitations; however, it is relatively labour-intensive, expensive, and dependent on the availability of high-quality reference sequences in the GenBank database and the selection of the appropriate gene fragment [24–26]. MALDI-TOF MS has been shown to be a fast, reliable and accurate method of identifying freshwater gastropod species, including intermediate hosts of schistosomiasis [7,12,21], and effective for tracing the geographic origin of freshwater snail species [6].

In Gabon, three species of schistosomes are found: *S. mansoni*, *S. intercalatum/guineensis*, which causes intestinal schistosomiasis, and *S. haematobium*, which causes urinary schistosomiasis [27–29]. However, malacological data are scarce in the country. The few data available on snails as intermediate hosts of schistosomiasis have been collected in the provinces of Estuaire and Moyen-Ogooué [29,30]. These morphological studies reported the presence of *Bu. globosus*, *Bu. forskalii* and *Bu. truncatus* in the surveyed freshwater bodies in the two provinces, but not that of *Biomphalaria sp*. Only the study carried out in Lambaréné by Dejon et al., provided evidence of cercarial excretion by *Bu. truncatus* snails, thus identifying this species as a potential intermediate host of *S. haematobium* in Gabon [29]. Furthermore, no significant association has been demonstrated between schistosomiasis transmission and the presence of *Bu. globosus* and *Bu. forskalii* snails [27–29]. However, *Bu. forskalii* and *Bu. globosus* have played a crucial role in the spread of schistosomiasis in Africa, and their presence in various parts of the country suggests a risk of pathogen transmission [12–14]. Therefore, it is therefore mandatory to study these snails and precisely describe what role they might play in the transmission of schistosomiasis in Gabon. Research has shown that *Bu. forskalii* can transmit both *Schistosoma intercalatum* and *S. guineensis*. Consequently, it is considered a potential intermediate host for these trematodes in Gabon and other regions [29]. Furthermore*, Bu. forskalii* has recently been identified as a potential intermediate host of *S. haematobium in Senegal* [12]. Nevertheless, intermediate host snails for schistosomiasis remain poorly studied and identified in several endemic areas of Gabon. There is no data on the potential presence of *Schistosoma* spp. in intermediate hosts in the semi-urban and rural areas of the Haut-Ogooué and Ogooué-Lolo provinces, where schistosomiasis has been reported [31]. Furthermore, the seasonal and geographical distribution of snails in Gabon is unclear. The aim of this study was to assess the spatial and temporal distribution of freshwater snails in rural and semi-urban areas of Gabon, to identify snail species using morphological, molecular, and protein-based (MALDI-TOF MS) approaches, and to detect *Schistosoma* spp. infections in snails through molecular techniques.

## METHODS

### STUDY AREAS

Freshwater snail surveys were conducted in the semi-urban areas of Koulamoutou and Lastoursville, and in the rural areas of Ndekabalendji, Okondja, Ondili, and Ayanabo, located in the Haut-Ogooué and Ogooué-Lolo provinces of southeastern Gabon (Figure 1). These regions are potentially endemic to schistosomiasis [32,33]. Local populations use watercourses near these localities, which are favourable to snails that act as intermediate hosts for *Schistosoma* spp. The region’s vegetation consists of tropical forests. The climate is hot and humid, with four distinct seasons: a long rainy season from mid-February to May, a long dry season from June to September, a short rainy season from October to mid-December and a short dry season from mid-December to mid-February. As part of this study, an in-depth analysis was carried out in six study areas to identify sites of contact between humans and water. These sites, which were identified as potential foci of schistosomiasis, were given particular attention. An in-depth analysis of the sites revealed significant vegetation cover. The riverbed in particular had a variety of substrates, such as sand, mud and pebbles, as well as combinations of these.

**Figure 1:**
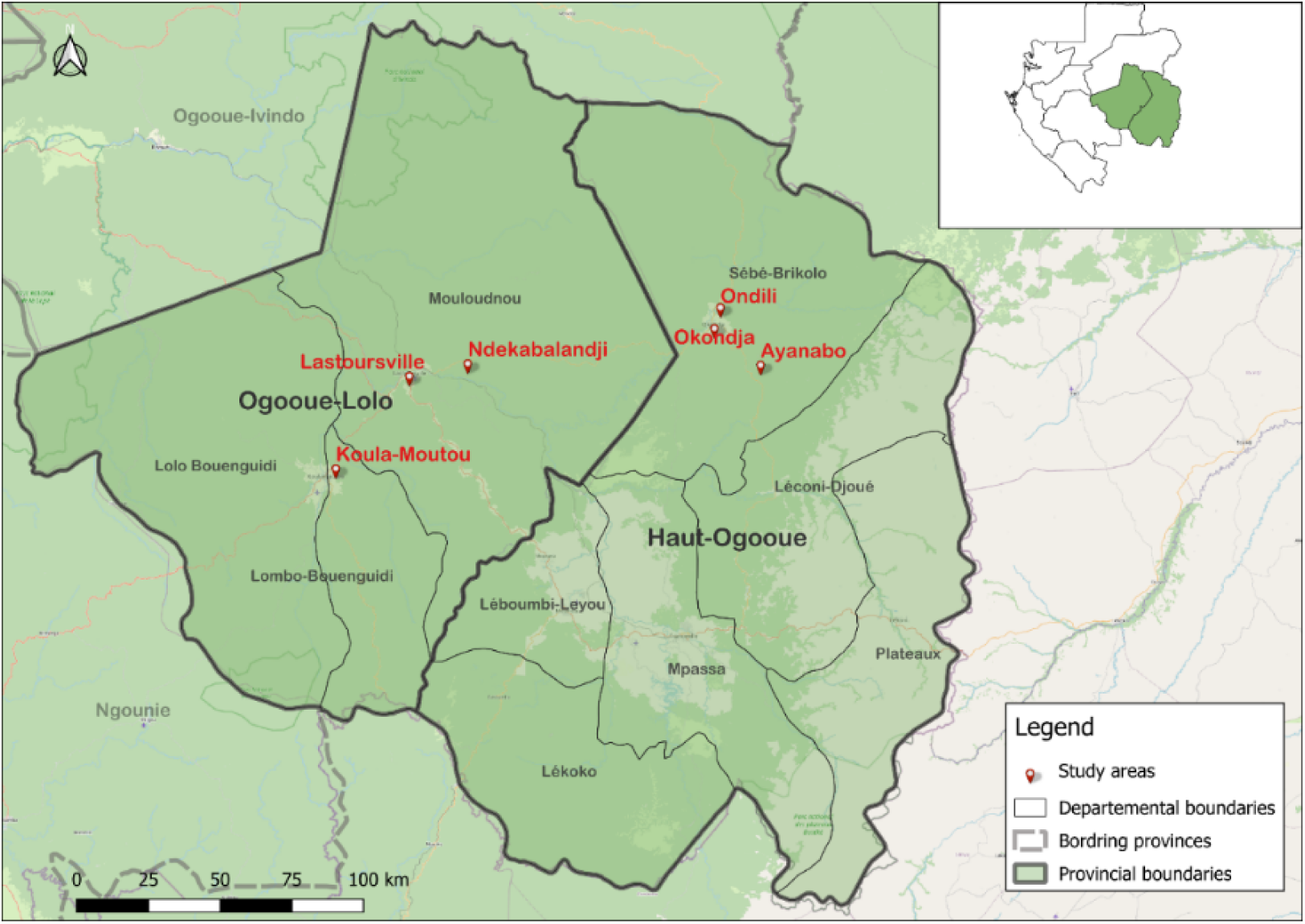
Map showing the study sites in the provinces of Ogooue-Lolo and Haut-Oguoué in Gabon.

#### Snail collection, morphological identification and examination of cercarial infection

Snails were collected once during three of the four seasons in Gabon. Thirty-seven survey sites were carefully selected based on their proximity to schools and dwellings, and to areas where human activity was observed for purposes such as recreation, domestic use, animal watering, agriculture and fishing. Two experienced and well-trained field collectors sampled the freshwater snails using long- and short-handled dip nets, depending on the water depth, as well as tongs. The snails were collected manually for 15 minutes. This involved digging along approximately 15 meters of riverbank, shaking out the vegetation, and turning over any objects left at the water’s edge. The snails were placed in plastic containers containing water and vegetation from the same habitat, then transported to the processing site. The snails were identified morphologically using a Zeiss Axio Zoom.V16 microscope (Zeiss, Marly-le-Roi, France) and standard identification keys [34]. Non-intermediate host snails of schistosomiasis were counted, recorded and returned to their respective sites, except for *Gyraulus costulatus*, which was present in large numbers in the waterways around Lastoursville railway station.

The intermediate host snails of schistosomiasis, as well as *G. costulatus*, were examined using the cercarial excretion method. The snails were placed individually in vials containing pure water and exposed to either sunlight or artificial light for between one and four hours, to induce excretion. The fluid from the vials was moved to a petri dish and observed under a stereomicroscope using an identification key [35,36] to identify the cercariae morphologically. After exposure, the snails were frozen for proteomic and molecular analysis. Images of the collection sites, the snails collected and the released cercariae can be found in Supplementary Figure S1. The snails were transferred to the Institut Hospitalo-Universitaire (IHU) in Marseille for MALDI-TOF MS and molecular analyses, under import permit no. ER69_2024.

#### Processing of snail specimens

The frozen snails were carefully extractedfrom their shells using soft forceps, rinsed with distilled water, and gently blotted dry using sterile filter paper. Each snail was then dissected using a sterile surgical blade undera binocular magnifying glass. The foot tissue was excised and placed in a 1.5 ml Eppendorf tube for MALDI-TOF MS analysis. The remainder of the snail, excluding the foot, was used for molecular biology analysis.

#### DNA extraction and molecular identification of snails

DNA-based identification was performed on a subsetof randomly selected specimens from each morphologically identified snail species. DNA was extracte from each snail’s tissue (excluding the foot) using the Machery Nagel ‘NucléoMag’ kit following the manufacturer’s instructions. The tissue samples were incubated overnight at 56 °C in 180 µl of NPL1 lysis buffer and 20 µl of proteinase K, until the cells were lysed and the tissue samples were homogenised. DNA extraction was subsequently carried out using the KingFisher™ Flex automated extraction system (Thermo Scientific™). Final elution was performed in 100 µl of elution buffer and the DNA was stored at −20°C for future use.

Standard PCR was performed on the DNA samples using an automated DNA thermal cycler (Applied Biosystems 2720, Foster City, USA). The targeted regions were the *18S rRNA* gene, for *Bu. truncatus* and *Bu. forskalii*, the *second internal transcribed spacer of the rRNA gene* (*ITS2*) for *Bu. truncatus* and the mitochondrial subunit I of *cytochrome oxidase* (*COI*) for *Bu. forskalii* and *G. costulatus*, which were amplified as previously described [7,12,37–39]. DNA from *Bu. truncatus*, which had previously been sequenced in the laboratory, was used as a positive control, and the PCR reaction mixture was used as a negative control. The primer sequences used are given in **Table S1**.

The amplification protocol consisted of an initial denaturation at 95°C for 15 min, followed by 40 cycles (45 cycles for *ITS* and 39 cycles for *18S*) at 95°C for 30 s, at 40°C for 30 s (at 45°C for 30 s for 18S and 42°C for 30 s for ITS), 72°C for 1 min (1 min for 18S) and a final step at 72°C for 7 min [37–39]. The PCR product was then migrated for 25 min at 180 V on a 1.5% agarose gel containing SYBR Safe dye, and the result was read using a Gel Doc system (Bio-Rad, Hercules, USA). The PCR products obtained were sequenced after purification using NucleoFast^TM^ 96 PCR (Macherey-Nagel, Düren, Germany), as previously described (1, 12). The ChromasPro ver. 1.7.7 software (Technelysium Pty Ltd, Tewantin, Australia) was used for sequence assembly and analysis. BLAST was performed to compare the obtained sequences with the reference sequences available in GenBank (http://blast.ncbi.nlm.nih.gov/). DNA sequences were randomly selected from each snail species to construct the phylogenetic tree. Initially, the sequences were aligned using BioEdit v7.2.5.0 software. They were then imported into MEGA X v10.2.6, where the phylogenetic trees were created by using the default maximum likelihood (ML) model. The number of nodes represented the percentage of bootstrap values obtained by repeating the analysis 100 times to generate a majority consensus tree, and only nodes with values of 80 or more were retained.

#### MALDI-TOF MS analysis

##### Sample processing and MALDI-TOF MS parameters

Foot tissue from each snail was homogenized with glass beads (Sigma, Lyon, France) in 40 µL of a solution containing of 70% (v/v) formic acid (Sigma) and 50% (v/v) acetonitrile (Fluka, Buchs, Switzerland) using a TissueLyser II apparatus (Qiagen, Hilden, Germany), as previously described [6,7,40]. Following brief centrifugation, 1 µL of the protein extract from each sample was spotted in quadruplicate onto a polished steel MALDI-TOF MS target plate (Bruker Daltonics, Wissembourg, France) [6,7,40]. After air drying at room temperature, each spot was overlaid with 1 µL of a 2.5% matrix solution, as described above [6,7,40].The target plate was then left to dry again at room temperature. Once dry, the target plate was introduced into the MALDI-TOF mass spectrometer (Bruker Daltonics, Germany). To control the loading process, the quality of the matrix solution and the performance of the MALDI-TOF MS apparatus, the matrix solution and laboratory-reared *anopheles gambiae* were loaded in duplicate onto each MALDI-TOF MS plate, serving as negative and positive controls, respectively.

The Microflex^LT^ MALDI-TOF mass spectrometer (Bruker Daltonics, Germany) was then used to generate protein mass profiles for each snail foot sample. Positive ions were detected in linear mode at a frequency of 50 Hz over a mass range of 2–20 kDa. Spectra were acquired using the default settings of the FlexControl v2.4 software (Bruker Daltonics) [41,42].

##### Spectral MS analysis creation of a reference spectra database

MS spectra obtained from snail foot tissues were visualized using FlexAnalysis v.3.3 (Bruker Daltonics, Germany), and subsequently exported to ClinProTools v.2.2 and MALDI-Biotyper v.3.0 (Bruker Daltonics, Germany) for further processing, including spectral smoothing, baseline subtraction, and intensity normalization (6,22). Spectra of insufficient quality defined as those with signal intensities below 3,000 arbitrary units (a.u.) or exhibiting excessive background noise were excluded from the analysis. Intra-species reproducibility and inter-species specificity of snail foot MS spectra were evaluated using clustering analysis (dendrogram) in MALDI-Biotyper v.3.0 and principal component analysis (PCA) in ClinProTools v.2.2. Principal spectra profiles (MSPs), generated from four replicate spots per sample, were used for clustering via the MSP dendrogram function in MALDI-Biotyper v.3.0 to assess spectral similarity among individuals of the same species. Additionally, PCA was performed using ClinProTools v.2.2 (default settings) to explore the distribution patterns of MS spectra between different snail species, as well as among populations from different geographical locations within the same species.

##### Creation of a reference spectra database and blind test

MS spectra from 12 snail specimens including four individuals from each species, whose morphological identification was confirmed by molecular analysis were used to update the internal MALDI-TOF MS reference database, which already contained spectra from several snail species (6,7). The remaining MS spectra that is, those not used to generate the reference database were compared againstthe updated MALDI-TOF MS reference spectra database using MALDI Biotyper v3.0 software. The degree of similarity between the query spectrum and the spectra in the database was evaluated using log score values (LSVs), which range from 0 to 3 depending on the level of matching. Identification was considered reliable when the LSV was ≥1.7 [7].

##### PCR detection of *Schistosoma* parasites

Real-time qPCR was used to detect *S. haematobium* infections in DNA extracts from snails. The 120 bp repetitive *Dra1* sequence of the *S. haematobium* complex was targeted using a hydrolysis probe developed by Cnops et al. [43] and primer pairs Sh1 and Sh2, as described by Hamburger et al. [44]. The primer sequences and probe used for qPCR are shown in **Table S1**. The reaction mixture consisted of 5 µL of DNA, 10 µL of Roche Master Mix (Roche, Basel, Switzerland), 3.5 µL of sterile distilled water, 0.5 µL of each primer (20 µM) and 0.5 µL of the probe (50 µM; Applied Biosystems, Foster City, CA, USA), giving a final volume of 20 µL. Amplification was performed using a CFX96 thermocycler (Bio-Rad, Marnes-la-Coquette, France). The thermocycler was programmed as follows: an initial denaturation step of 2 minutes at 95 °C, followed by a denaturation step of 3 minutes at 95 °C, then 40 cycles of 30 seconds at 95 °C and 1 minute at 60 °C, followed by holding at 4 °C indefinitely. The qPCR result was considered positive when the cycle threshold (Ct) value was less than 35. Negative (distilled water) and positive controls were used for each PCR.

To refine parasite identification, some qPCR-positive samples were sequenced using *ITS2* [45] and *COX1* [45,46] primers. The DNA obtained from each sample was used in the PCR reaction. The primer sequences used are given in **Table S1** [45,46]. The amplification protocol involved an initial denaturation step at 95 °C for 15 minutes, followed by 39 cycles of denaturation at 95 °C for 30 seconds, annealing at 58 °C (or 56 °C for *ITS2*) for one minute, and extension at 72 °C for one minute. This was followed by a final extension step at 72 °C for seven minutes. The same protocol used for snail identification was applied to the sequencing process.

##### Data analysis

The data were recorded in Microsoft Excel 2013 and analysed using R version 4.4.0. A chi-squared test was performed to compare prevalence rates. A p-value of less than 0.05 was considered significant. Overall snail abundance by species was calculated by dividing the number of snails of a given species by the total number of species collected. Seasonal abundance was calculated by dividing the number of snails of a given species by the total number of snails collected during that season. The data were presented in figures and tables. Geographical coordinates were collected using a Garmin GPSMAP 66i and then used to create maps. The data were imported into a Jupyter Notebook and processed using Python 3. The data was sorted using the Pandas library, and the maps were created using the Folium library.

## RESULTS

### Freshwater snail collections and morphological identification

A total of 3,722 freshwater snail specimens were collected from all the surveyed water points in Lastoursville (n = 3,352), Okondja (n = 94), Ayanabo (n = 176), Koulamoutou (n = 14) and Ndekabalandji (n = 86) (**Figure 2**). Based on morphological characteristics, the snails were identified as belonging to four genera: *Bulinus* (n =2226), *Gyraulus* (n =680), *Melania* (n =749), and *Bellamya* (n =67) (Table 1 and Table S2). Specimens from the genus *Bulinus* were further identified at the species level as *Bulinus forskalii* and *B. truncatus*, while specimens from the genus *Gyraulus* were identified as *Gyraulus costulatus* (Table 1).

**Figure. 2.**
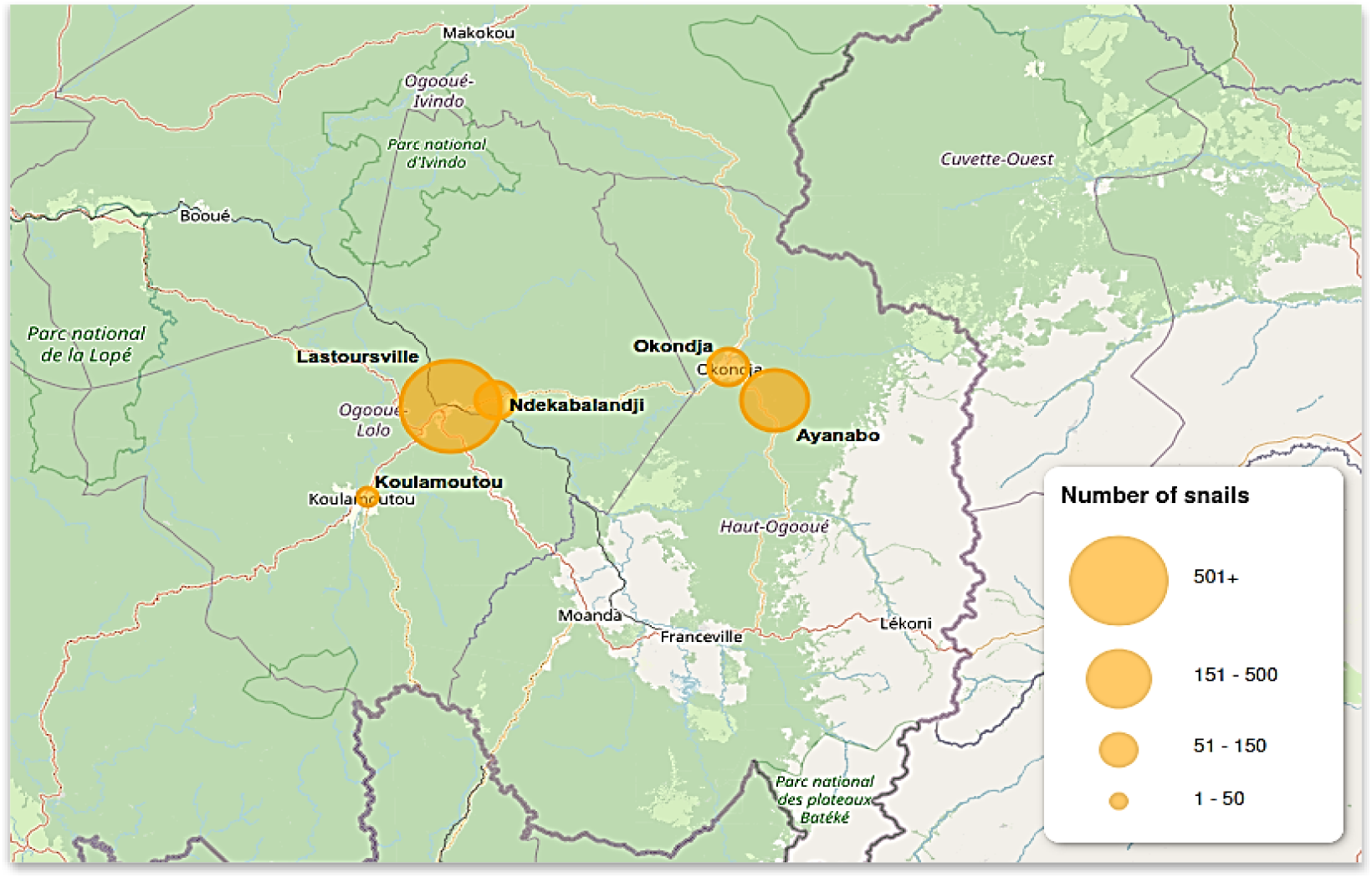
Overall distribution of snails collected by study sites. Distribution and abundance of *Bu. truncatus*, *Bu. forskalii* and *G. costulatus*.

**Table 1** shows the spatial distribution and abundance of the two potential intermediate host snail species of schistosomiasis (*Bu. forskalii* and *Bu. truncatus*) and *G. costulatus* (n = 2,906). Overall, *B. truncatus* snails were the most abundant (n = 1,845, A = 0.63), followed by *G. costulatus* (n = 680, A = 0.24) and *B. forskalii* (n = 381, A = 0.13) (p < 0.05).

**Table. 1.** Overall distribution and abundance of *Bulinus* and *Gyraulus* snail species by locality, based on morphology.

| LOCALITIES<br>(N collection sites) | *B. truncatus* | *B. forskalii* | *G. costulatus* | Total |
| --- | --- | --- | --- | --- |
| Lastoursville | 1834 (13) | 114 (5) | 610 (14) | 2558 |
| Koulamoutou | 0 | 3(3) | 11 (2) | 14 |
| Ndekabalandji | 9 (1) | 0 | 56 (1) | 65 |
| Okondja | 2 (1) | 89 (3) | 3 (1) | 94 |
| Ayanabo | 0 | 175 (3) | 0 | 175 |
| **Total (n)** | 1845 | 381 | 680 | 2906 |
| **Abundance** | 0.63 | 0.13 | 0.24 |  |

A detailed analysis of the data collected during the survey revealed that snails were observed at 28 of the 37 assessed collection points: 18 in Lastoursville, four in Okondja, three in Ayanabo, two in Koulamoutou, and one site in Ndekabalandji (Supplementary **Table S2** and **Figures 3A, 3B, and 3C**). No snail was found at any of the Ondili sites surveyed. Figure 3 provides a visual representation of the distribution of snail species, while the data is further broken down in the detailed table provided in the supplementary material (**Table S2**).

**Figure. 3.**
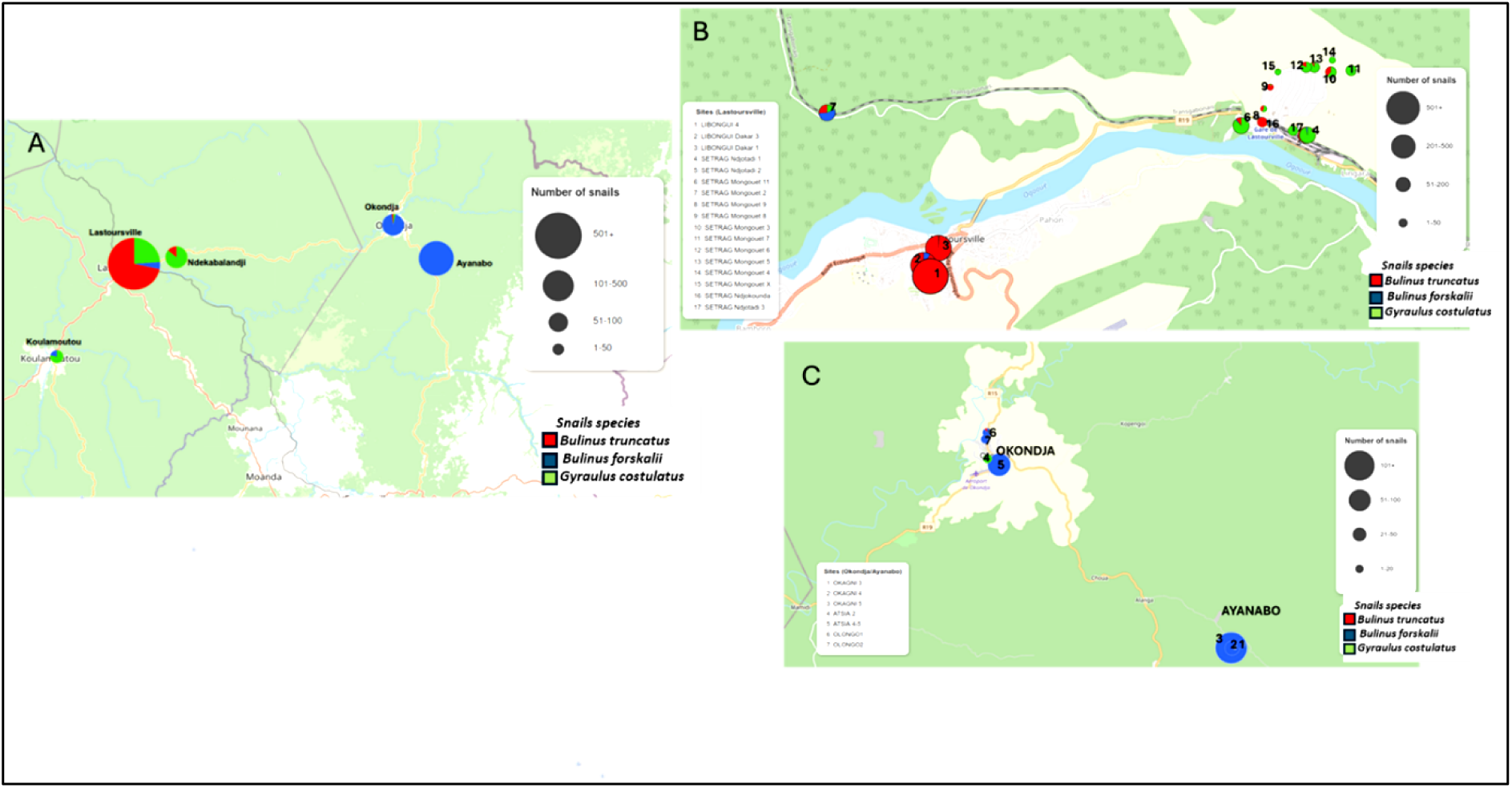
Spatial distribution of snail species collected by study sites. **A**-Map showing the species of snails collected in the five localities. **B-** Distribution of snail species in the Lastoursville sites. **C**-Distribution of snail species in the Ayana and Okondja sites.

### Snail abundance and seasonality

**Figure 4** shows the seasonal distribution of the snail species collected. The number of *Bu. truncatus*, *Bu. forskalii* and *G. costulatus* varied significantly according to season. Comparing the overall abundance of snails by season reveals that they are more abundant at the beginning (n = 1,660, A = 0.57) and end (n = 902, A = 0.3) of the long dry season than during the short dry season (n = 339, A = 0.11) and long rainy season (n = 5, A = 0.001), p < 0.05. At the beginning of the long dry season, 51.5% (865), 27.4% (455) and 20.5% (340) of the snails were *Bu. truncatus*, *G. costulatus* and *Bu. forskalii*, respectively. *Bu. truncatus* was the most widespread species (p < 0.05). Similarly, snails collected at the end of the long dry season and during the short dry season showed a significantly higher prevalence of *Bu. truncatus* than *G. costulatus* and *Bu. forskalii* (p < 0.05). In contrast, during the long rainy season, the proportions of the different snail species collected did not differ significantly and *G. costulatus* was absent.

**Figure. 4.**
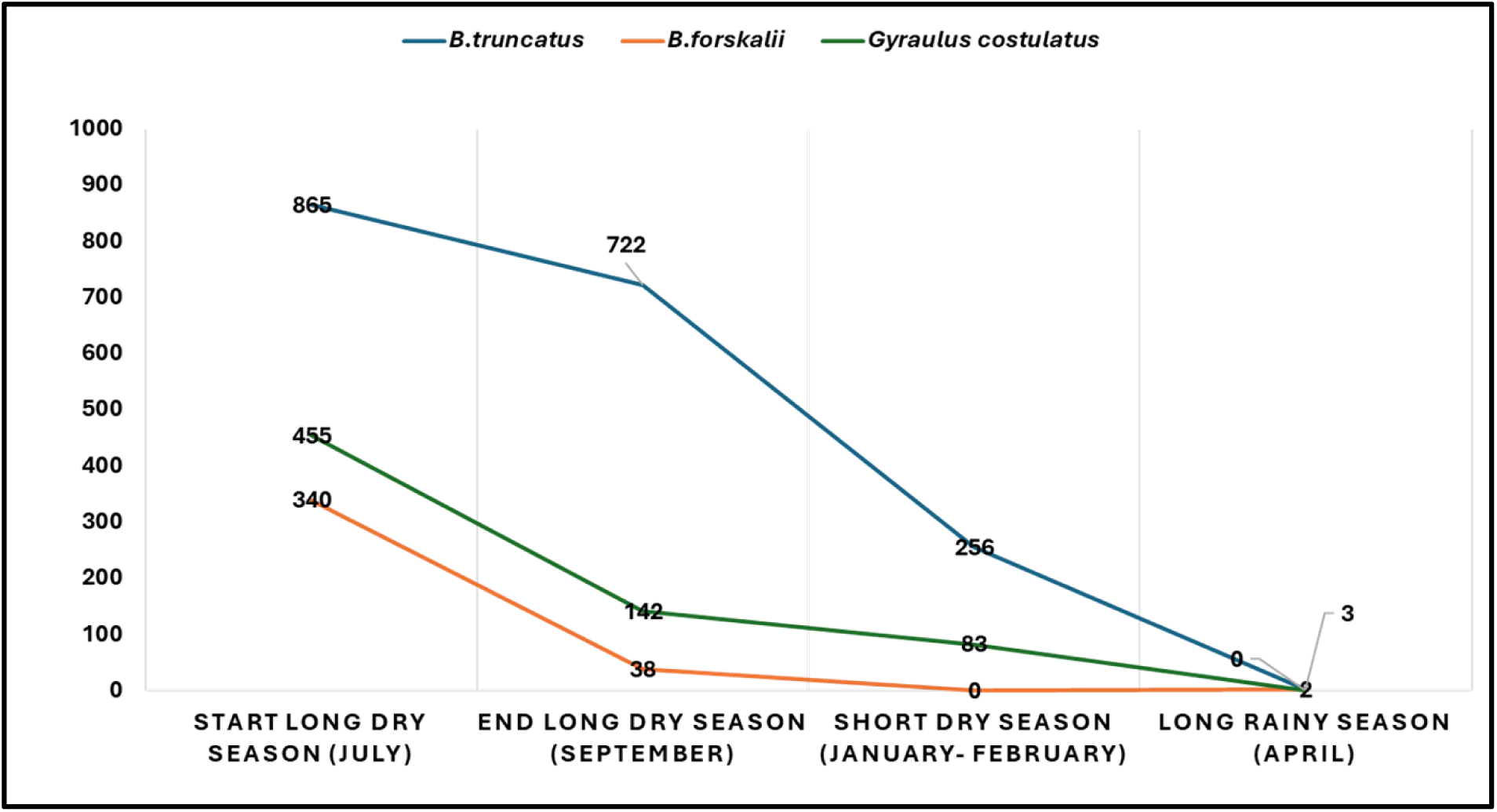
Seasonal distribution of snails.

### Cercarial emission test

Both species of intermediate host snail tested released cercariae of *Schistosoma* spp. Intriguingly, cercariae were discharged by *G. costulatus*, which was unanticipated. A total of 426 snails were found infested, giving an overall 14.7% infestation rate (426 out of 2,890). Of these, 308 (16.7%) were *Bu. truncatus*. *Bu. forskalii* had an 11.2% infestation rate (43 out of 381), while *G. costulatus* had a 13% infestation rate (85 out of 680). Cercariae excreting *Bu. truncatus* were collected only at the Lastoursville site. Infested *Bu. forskalii* were collected at Lastoursville, Ayanabo and Okondja. Infested *G. costulatus* were collected at Lastoursville and Ndekabalandji. No infested snails were collected at Koulamoutou (**Table S2**).

### DNA sequence-based snail identification

A total of 68 morphologically identified snails (35 *Bu. forskalii*, 26 *Bu. truncatus* and 7 *G. costulatus*) were subjected to standard PCR and sequencing. Twelve of these snail specimens were used to update the internal MALDI-TOF MS database (four of each species). Fourteen were identified morphologically, then by MALDI-TOF MS, and were positive for schistosomiasis. Fifty presented an LSV <1.7 when tested blindly against the updated reference database. The success rate for *18S rRNA* gene-based sequencing was 100% (61/61) for *Bu. forskalii* and *Bu. truncatus*, with at least one sequence obtained for each species. *Bu. truncatus* specimens were re-sequenced targeting the *ITS* gene, while *G. costulatus* and *Bu. forskalii* specimens were sequenced targeting the *COI* gene. The sequencing success rate for these two genes was 100%. BLAST analysis of the *Bu. forskalii* and *G. costulatus COI* sequences showed identities ranging from 98 to 100% with GenBank reference sequences AM286308, ON077052 and OP811018 respectively (see Table 3 for details). Comparing the 18S gene sequences obtained from *Bu. forskalii* and *Bu. truncatus* with the GenBank reference sequences KJ157347.1 and HM756320.1 revealed identity percentages ranging from 97% to 100%. However, variation was observed at the species level for the 18S gene. In fact, this marker did not enable precise discrimination between species belonging to the Africanus and Tropicus groups. Furthermore, 100% intraspecific identity was observed for all *Bu. truncatus* snail sequences obtained with the *ITS* gene, as well as with GenBank sequences (KJ157504.1 and ON553184.1) (**Table 2**).

**Table. 2.** Results of the DNA-based identification of the snails.

| Morphological identification | Number of snails tested<br>By ITS2 | BLAST best-match GB<br>Accession No | Number of snails tested<br>By COI | BLAST best-match GB<br>Accession No | Number of snail tested<br>By 18S | BLAST best-match GB<br>Accession No |
| --- | --- | --- | --- | --- | --- | --- |
| *Bulinus truncatus* | 26 | KJ157504.1<br>ON553184.1 | NA | NA | 26 | KJ157347.1 |
| *Bulinus forskalii* | NA | NA | 35 | AM286308.2 | 35 | HM756320.1 |
| *Gyraulus costulatus* | NA | -NA | 7 | OP811018.1 |  | NA |
| **Total** | **26** | – | **42** |  | **61** | - |

**Table. 3.** Results of the MALDI-TOF MS identification of the snails.

| Morphological identification | Number of snail tested | Good quality spectra | Spectra added to the data base (N) | Blind test | Good identification (%) | Log Score Values range |
| --- | --- | --- | --- | --- | --- | --- |
| *Bulinus truncatus* | 430 | 305 | 4 | 301 | 289 (96%) | 1,71-2,8 |
| *Bulinus forskalii* | 194 | 151 | 4 | 147 | 147 (100%) | 1,71-2,5 |
| *Gyraulus costulatus* | 167 | 162 | 4 | 158 | 158 (100%) | 1,71-2,5 |
| **Total** | **791** | **618** | **12** | **606** | - | - |

The representative sequences (*COI, 18S rRNA* and *ITS2*) of *Bu. forskalii*, *Bu. truncatus* and *G. costulatus* have been deposited in the GenBank nucleotide database under the accession numbers [PV392216-PV392223](https://www.ncbi.nlm.nih.gov/nuccore/?term=PV392216:PV392223%5baccn%5d) (*COI*), [PV399981-PV399987](https://www.ncbi.nlm.nih.gov/nuccore/?term=PV399981:PV399987%5baccn%5d) (*18S*) and [PV413072-PV413076](https://www.ncbi.nlm.nih.gov/nuccore/?term=PV413072:PV413080%5baccn%5d) (*ITS2*). These sequences are available in FASTA format in Supplementary File S3. The phylogenetic tree below clearly shows the clustering of snail species from Gabon with those from Nigeria, Cameroon, Niger and Senegal (**Figure 5**).

**Figure. 5.**
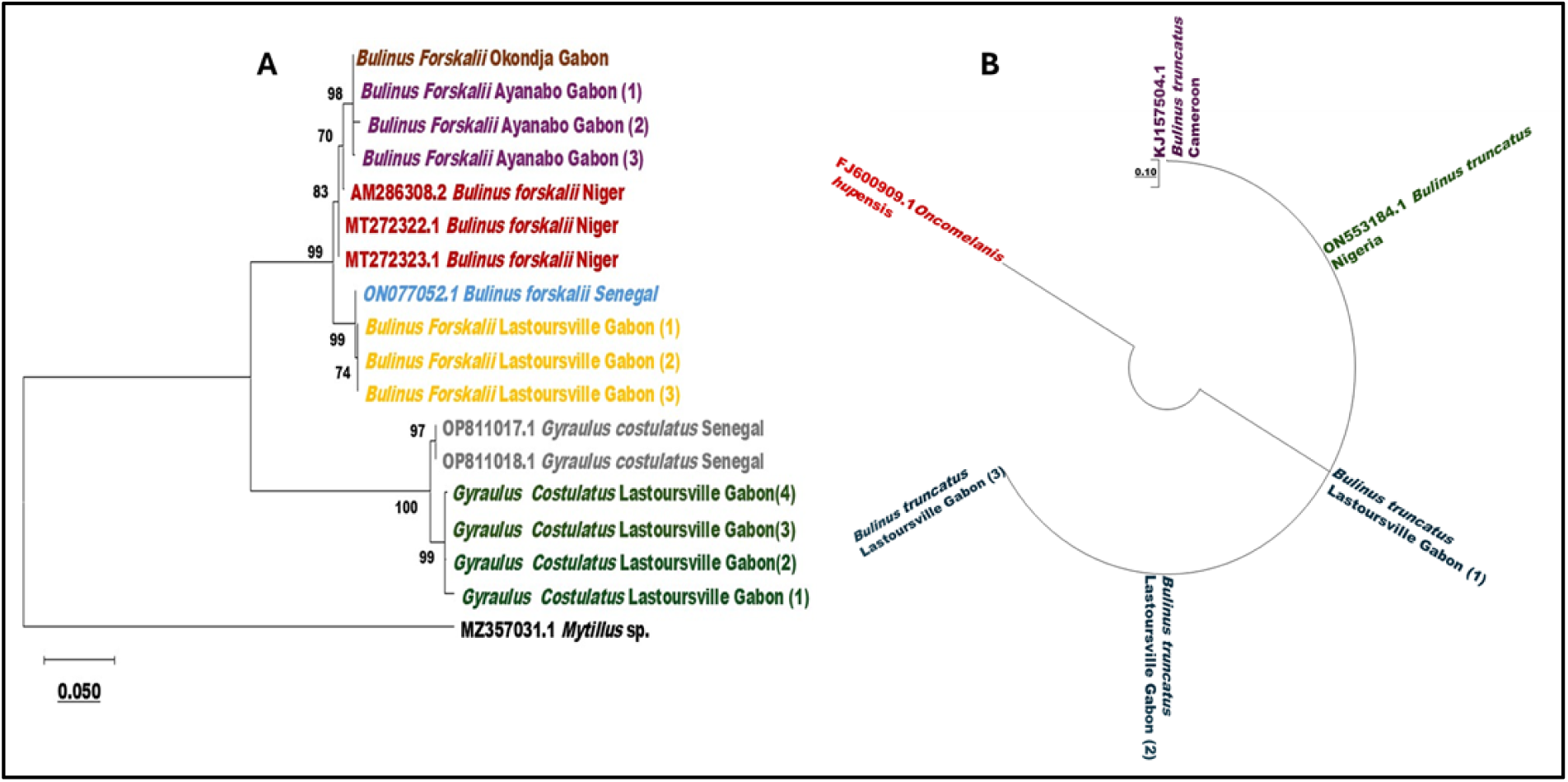
Phylogenetic tree showing the position of the snail species sequenced. A- Trees constructed using COI gene sequences. B- Tree built with ITS2 gene sequences. The tree was constructed using the maximum likelihood method based on the Kimura.

### MS spectra analysis

A total of 791 snails including 301 *Bulinus truncatus*, 158 *B. forskalii*, and 147 *Gyraulus costulatus* were analyzed by MALDI-TOF MS. Visualization of the spectra using FlexAnalysis v.3.3 software showed that 78.2% (n = 618) of the specimens produced high-quality spectra, defined by a maximum intensity >3,000 arbitrary units (a.u.) and the absence of background noise (Table xx). Visual comparison of the overlaid MS spectra for the three species revealed high intra-species reproducibility and clear inter-species specificity. Figure 6A). To further assess the reproducibility and specificity of the MS spectra among different snail species, clustering and principal component analysis (PCA) were performed. The dendrogram generated from spectra of four specimens per species showed that individuals belonging to the same species clustered together on the same branches (Figure 6B). Similarly, PCA demonstrated consistent grouping, with specimens of the same species forming distinct clusters (Figures 6C). Interestingly, PCA analyses performed on *Bulinus truncatus* and *B. forskalii* specimens collected from different geographical locations revealed clear clustering patterns according to collection site, as shown in Figures S4.A and S4.B for *B. truncatus* and *B. forskalii*, respectively.

**Figure 6.**
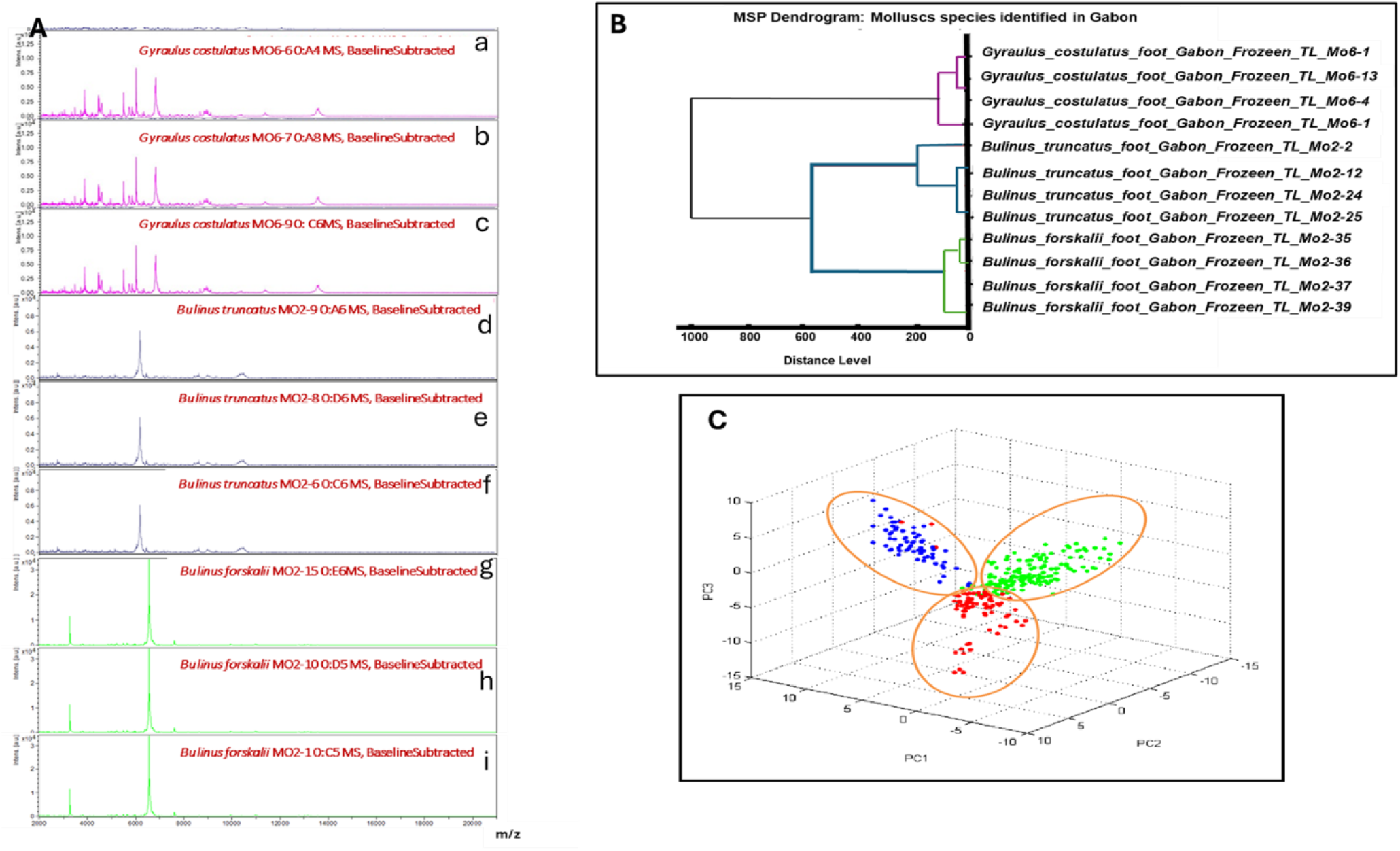
MALDI-TOF MS spectra acquired from snail foot samples. (A)- Representative MALDITOF MS profile of *Gyraulus costulatus* (a, b, c), *Bulinus truncatus* (d, e, f), and *Bulinus forskalii* (g, h, i), obtained with the FlexAnalysis v.3.3 software. (B)- Hierarchical clustering dendrogram of *Bulinus truncatus*, *Gyraulus costulatus*, *Bulinus forskalii*, obtained from the software Biotyper v.3.0. (C)- Graphical representation (PCA), determining the classification of the LSV of the species present (blue colour = *Gyraulus costulatus*, red colour = *Bulinus forskalii*, green colour = *Bulinus truncatus*), using the software ClinProTools v.2.2.

### Blind test for MALDI-TOF MS identification of snail species

From the high-quality spectra of 618 specimens, those of 12 individuals four from each species with morphological identifications confirmed by molecular biology were selected to update the MALDI-TOF MS database. The spectra of the remaining 606 specimens were then blindly tested against this updated database. All *G. costulatus* (n = 158) and *Bu. forskalii* (n = 147) specimens were successfully identified, with log score values (LSVs) ranging from 1.71 to 2.5. In contrast, 96.01% (289 out of 301) of *Bu. truncatus* spectra were correctly identified, with LSVs ranging from 1.71 to 2.8 (Table 3).

### qPCR detection of *Schistosoma* parasites

qPCR targeting the *DraI* gene of the *S. haematobium* complex was performed on 606 snail specimens that had been identified using MALDI-TOF MS. The results revealed *Schistosoma* spp. in 107 specimens (17.66%), including 83 *Bu. truncatus* (28.7%), 11 *Bu. forskalii* (7.5%) and 13 *G. costulatus* (8.3%), with cycle threshold (Ct) values ranging from 13 to 34.9. *COX1* gene amplification yielded a 543-base-pair fragment for *S. haematobium* in four *G. costulatus* and five *Bu. forskalii* species. All 83 *Bu. truncatus* specimens were positive for *S. haematobium*. To confirm the exact species of *Schistosoma,* present in the three snail species, eight *B. truncatus*, eleven *B. forskalii* and thirteen *G. costulatus* specimens were randomly selected for further DNA sequence analysis. Sequencing of the *ITS2* and *COX1* regions definitively identified *S. haematobium* in *Bu. truncatus* (n = 8), *G. costulatus* (n = 2) and *Bu. forskalii* (n = 3) (**Table 4**).

**Table. 4.**
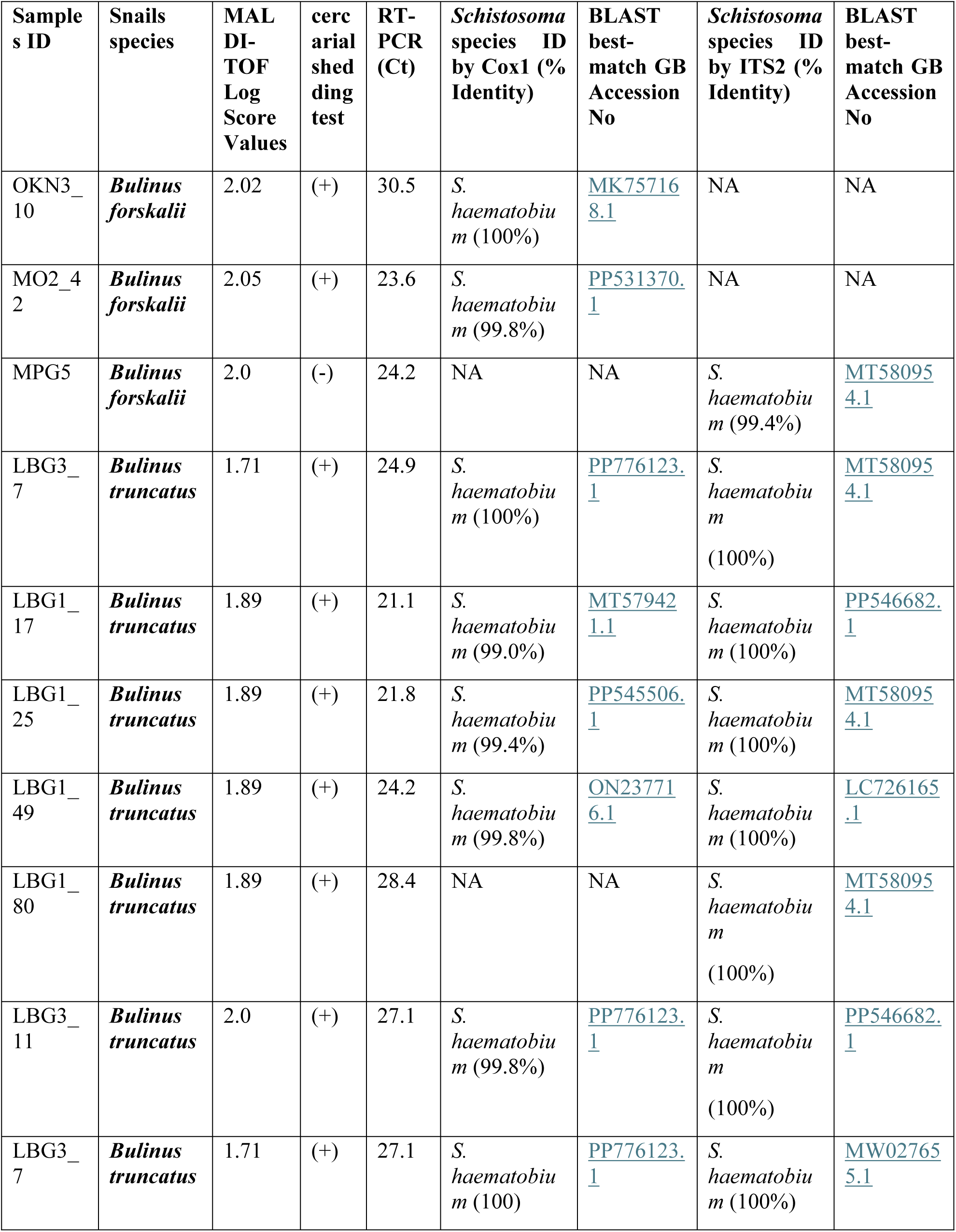
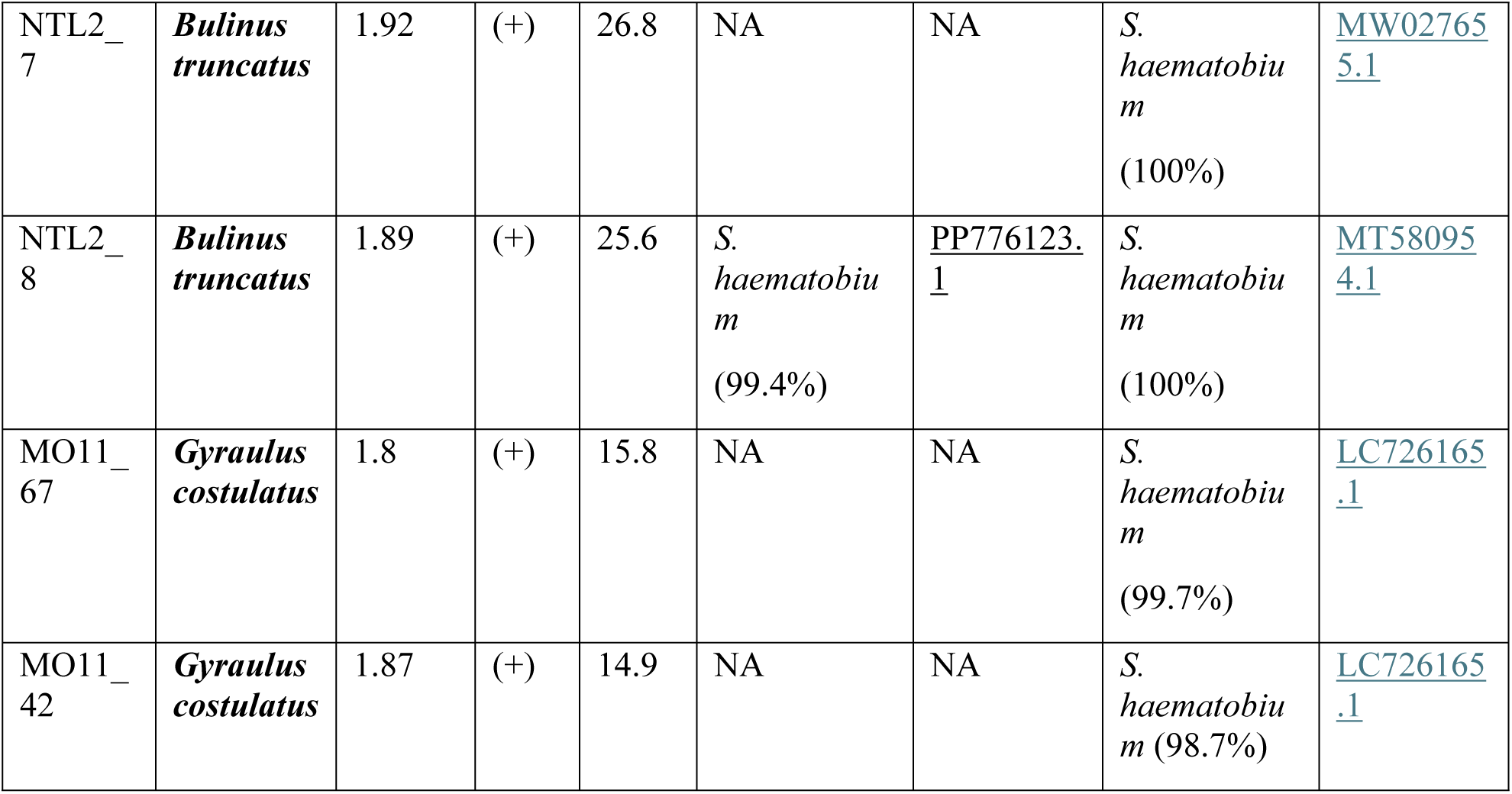
MALDI-TOF MS identification of the snail positive for *S*. *hæmatobium* complex PCR, and COX1 and ITS2 DNA sequence-based identification of the *Schistosoma* sp., parasites.

| Sample<br>s ID | Snails<br>species | MAL<br>DI-TOF<br>Log<br>Score<br>Values | cerc<br>arial<br>shed<br>ding<br>test | RT-PCR<br>(Ct) | *Schistosoma*<br>species ID<br>by Cox1 (%<br>Identity) | BLAST<br>best-match GB<br>Accession<br>No | *Schistosoma*<br>species ID<br>by ITS2 (%<br>Identity) | BLAST<br>best-match GB<br>Accession<br>No |
| --- | --- | --- | --- | --- | --- | --- | --- | --- |
| OKN3_<br>10 | *Bulinus*<br>*forskali* | 2.02 | (+) | 30.5 | *S.*<br>*haematobiu*<br>*m* (100%) | MK75716<br>8.1 | NA | NA |
| MO2_<br>4<br>2 | *Bulinus*<br>*forskali* | 2.05 | (+) | 23.6 | *S.*<br>*haematobiu*<br>*m* (99.8%) | PP531370.<br>1 | NA | NA |
| MPG5 | *Bulinus*<br>*forskali* | 2.0 | (-) | 24.2 | NA | NA | *S.*<br>*haematobiu*<br>*m* (99.4%) | MT58095<br>4.1 |
| LBG3_<br>7 | *Bulinus*<br>*truncatus* | 1.71 | (+) | 24.9 | *S.*<br>*haematobiu*<br>*m* (100%) | PP776123.<br>1 | *S.*<br>*haematobiu*<br>*m*<br>(100%) | MT58095<br>4.1 |
| LBG1_<br>17 | *Bulinus*<br>*truncatus* | 1.89 | (+) | 21.1 | *S.*<br>*haematobiu*<br>*m* (99.0%) | MT57942<br>1.1 | *S.*<br>*haematobiu*<br>*m* (100%) | PP546682.<br>1 |
| LBG1_<br>25 | *Bulinus*<br>*truncatus* | 1.89 | (+) | 21.8 | *S.*<br>*haematobiu*<br>*m* (99.4%) | PP545506.<br>1 | *S.*<br>*haematobiu*<br>*m* (100%) | MT58095<br>4.1 |
| LBG1_<br>49 | *Bulinus*<br>*truncatus* | 1.89 | (+) | 24.2 | *S.*<br>*haematobiu*<br>*m* (99.8%) | ON23771<br>6.1 | *S.*<br>*haematobiu*<br>*m* (100%) | LC726165<br>.1 |
| LBG1_<br>80 | *Bulinus*<br>*truncatus* | 1.89 | (+) | 28.4 | NA | NA | *S.*<br>*haematobiu*<br>*m*<br>(100%) | MT58095<br>4.1 |
| LBG3_<br>11 | *Bulinus*<br>*truncatus* | 2.0 | (+) | 27.1 | *S.*<br>*haematobiu*<br>*m* (99.8%) | PP776123.<br>1 | *S.*<br>*haematobiu*<br>*m*<br>(100%) | PP546682.<br>1 |
| LBG3_<br>7 | *Bulinus*<br>*truncatus* | 1.71 | (+) | 27.1 | *S.*<br>*haematobiu*<br>*m* (100) | PP776123.<br>1 | *S.*<br>*haematobiu*<br>*m* (100%) | MW02765<br>5.1 |
| NTL2_<br>7 | ***Bulinus truncatus*** | 1.92 | (+) | 26.8 | NA | NA | *S. haematobium*<br>(100%) | MW02765<br>5.1 |
| NTL2_<br>8 | ***Bulinus truncatus*** | 1.89 | (+) | 25.6 | *S. haematobium*<br>(99.4%) | PP776123.1 | *S. haematobium*<br>(100%) | MT58095<br>4.1 |
| MO11_<br>67 | ***Gyraulus costulatus*** | 1.8 | (+) | 15.8 | NA | NA | *S. haematobium*<br>(99.7%) | LC726165<br>.1 |
| MO11_<br>42 | ***Gyraulus costulatus*** | 1.87 | (+) | 14.9 | NA | NA | *S. haematobium*<br>(98.7%) | LC726165<br>.1 |

The representative sequences (ITS2 and COI) of *Schistosoma haematobium* have been deposited in the GenBank nucleotide database under the accession numbers [PV413076-PV413080](https://www.ncbi.nlm.nih.gov/nuccore/?term=PV413072:PV413080%5baccn%5d) and [PV796177-PV796179](https://www.ncbi.nlm.nih.gov/nuccore/?term=PV796177:PV796179%5baccn%5d). These sequences are available in FASTA format in Supplementary File S3. The phylogenetic tree below clearly shows the clustering of Schistosoma haematobium from Gabon with those from Niger, France, and Senegal (**Figure.S4**).

All *Bu. truncatus* and *G. costulatus* snails that tested positive for *S. haematobium* originated from Lastoursville. The *Bu. forskalii* that tested positive for *S. haematobium* were collected in Ayanabo, Lastoursville and Ndekbalandji (**Figure.S4**).

## Discussion

This study is the first to accurately identify frozen specimens of different snail species in Gabon. This is the first study in Gabon to use the state-of-the-art MALDI-TOF MS method to identify the freshwater snails which transmit schistosomiasis. These results add to the many epidemiological studies in schistosome-endemic areas. To plan and implement effective prevention and control strategies, it is necessary to understand the dynamics of local transmission and to be aware of the malacological fauna of snail intermediate hosts of schistosomiasis [10]. While snail species inventories in Africa have traditionally relied on morphological identification, the recent adoption of MALDI-TOF MS as a powerful, inexpensive, rapid identification tool for arthropods, marine bivalves, and certain gastropods (including schistosomiasis intermediate hosts) has transformed medical entomology and malacology [6,7,26,47,48]. In Gabon, no study has yet been conducted on identifying snails using MALDI-TOF MS. Our study supplements the limited malacological data available in Gabon and presents the first such data from the Ogooué-Lolo and Haut-Ogooué provinces obtained using innovative techniques. Additionally, this study demonstrated that seasonal changes impact snail abundance, as previously reported [49].

The four genera of freshwater snail identified based on morphology in the water bodies of the five localities studied in Gabon had previously been recorded in Gabon [27,29]. Consistent with previous studies observing cohabitation between *Bulinus* and other snails [29,50,51], we found that some sites were home to up to three snail genera, indicating the coexistence of different species within the same biotope.

The snails collected were identified as *Bu. truncatus*, *Bu. forskalii*, *G. costulatus*, *Melania* sp. and *Bellamya* sp. The most abundant species were *Bu. truncatus*, *Bu. forskalii* and *G. costulatus*; the former two are implicated in the transmission of schistosomiasis. *G. costulatus* is an invasive species of snail that has already been identified and described in some African countries. It has extensively colonised bodies of water at 18 sites in Lastoursville (n = 14), Koulamoutou (n = 2), Okondja (n = 1) and Ndekabalandji (n = 1) and is widely distributed and abundant in Lastoursville. However, its role as an intermediate host of trematodes has never been documented. Both *Bu. truncatus* and *Bu. forskalii* were widely distributed across most of the surveyed water points. B. truncatus snails colonised 15 sites: 13 in Lastoursville, one in Okondja and one in Ndekabalandji, where they were particularly abundant. Bu. forskalii snails were present at 12 sites: five in Lastoursville, three in Okondja, one in Koulamoutou and three in Ayanabo. The highest abundance was noted in Lastoursville and Ayanabo. These data are consistent with those reported by Déjon et al. in rural and semi-urban areas of the Moyen-Ogooué province [29]. Furthermore, Mintsa Nguema et al.’s study reported the presence of *Bu. forskalii* and *Bu. globosus* in the estuary, but not *Bu. truncatus* [27]. Studies conducted in Cameroon and the Republic of the Congo have also identified *Bu. forskalii* and *Bu. truncatus* as intermediate hosts of schistosomiasis [35,52]. Similarly, several studies conducted in West Africa have described the presence of these two species, which are well known to be involved in the transmission of schistosomiasis in this region [7,12]. These data demonstrate the sub-regional distribution of *Bulinus* species and their widespread presence across Africa, which is facilitated by environmental conditions that support their development. Given the broad dissemination of these gastropods, the water sources used by populations in Lastoursville, Okondja, Ayanabo, Ondili and NdekabaLandji could be the main transmission sites for urogenital and intestinal schistosomiasis in Gabon. *S. haematobium* and *S. guineensis* have already been reported in some of the localities studied [31,53].

We found that there was a higher abundance of snails during the dry season, with significant reductions occurring during the rainy season. These population dynamics are probably the result of several abiotic factors acting together. For instance, snail displacement is restricted during the wet months due to increased water depth and flow, which reduces their numbers. Several studies [50,54,55] have also demonstrated a negative correlation between snail distribution and abundance, and rainfall. In line with these studies’ conclusions, we confirmed that the dry season is the best time to collect snails.

In this study, DNA-based analyses were employed to corroborate the morphological identification of nearly all snail species. The *mitochondrial cytochrome oxidase subunit I* (*COI*) gene, the *internal transcribed spacer 2* (*ITS2*) gene and *the 18S ribosomal RNA* (*rRNA*) gene were used to identify the necessary snail species for enriching the MALDI-TOF MS database and to distinguish between species that could not be identified by MALDI-TOF MS alone. Previous studies have reported that the COI gene is an effective marker for discriminating between *Bulinus* species due to the high degree of sequence divergence present in this gene. Our study confirmed the identities of *Bu. forskalii* and *G. costulatus*, thus demonstrating the importance of targeting the *COI* gene for snail species identification. These data could unambiguously discriminate between species within the same group using *COI* [7]. In our study, the *ITS2* gene was found to be an effective marker for identifying *Bu. truncatus*. Indeed, all morphologically identified *Bu. truncatus* species showed 100% identity with the GenBank KJ157504.1 sequence. However, querying the GenBank database with *18S rRNA* gene sequences was unable to classify the sequences according to species. The *18S rRNA* gene did not permit clear differentiation between snails belonging to the *Bu. truncatus*/Tropicus group and the *Bu. forskalii/africanus* group. These results are comparable to those found in diversity studies, which have shown that *18S* gene fragments exhibit lower variation than the more variable *ITS* fragment. This suggests that the *ITS* gene is better able to distinguish between species within the same group than the 18S gene, which is unable to discriminate between them [56,57].

The cercariae emission test revealed that the collected Bu. truncatus snails emitted Schistosoma spp. cercariae. This finding is consistent with a study by Déjon et al. in Lambaréné, which suggests that *Bu. truncatus* is the intermediate host of urinary schistosomiasis in the region. However, the present study provides evidence that *Bu. forskalii* snails excrete cercariae, contrasting with previous study data that found no infested *Bu. forskalii* [29]. However, complementary analyses based on RT-PCR (DraI) and sequencing (ITS and COX1) confirmed the presence of *S. haematobium* in *Bu. forskalii* and *Bu. truncatus* samples. This establishes them as intermediate hosts of *S. haematobium* in Gabon. The detection of *S. haematobium* in *Bu. forskalii* corroborates the findings of a study of specimens collected in Senegal which also reported the presence of *S. haematobium* in *Bu. forskalii*[12]. Nevertheless, studies conducted in Central Africa failed to detect *S. haematobium* in *Bu. forskalii*, despite earlier descriptions of this snail as an intermediate host of *S. guineensis/intercalatum* [58]. Our results also showed that *S. haematobium* could infest the invasive snail species *G. costulatus*, which was collected in Lastoursville. No case of schistosomiasis infestation in this snail has yet been reported. We hypothesise that *G. costulatus* could be involved in the transmission of schistosomiasis and therefore warrants special attention in future studies. To investigate this further, natural infestation tests would need to be carried out, with individual cercariae analysed for both *Bu. forskalii* and *G. costulatus*.

This study has two limitations. Firstly, the geographical scope of the study was limited to two provinces. Therefore, it should be extended to other areas of Gabon. Secondly, the emitted cercariae were not analysed, hindering the precise identification of the parasite.

## Conclusions

Our study provides the first MALDI-TOF MS identification of freshwater snails collected in Gabon, thereby adding to the existing malacological data for the country. Our findings confirm the presence of *Bu. forskalii* and *Bu. truncatus* in Gabon’s freshwater habitats, both of which are endemic to regions where schistosomiasis is prevalent. They also demonstrate the heterogeneous geographical distribution of these snails in Gabon. Furthermore, our study confirms that snail collection is more successful in the dry season than in the rainy season. Additionally, our results confirm the infestation of *Bu. forskalii* and *Bu. truncatus* by *S. haematobium*, suggesting that they act as intermediate hosts in Gabon. However, further study is required on the role of other snail species collected in Gabon’s freshwater bodies in the transmission of schistosomiasis, including the invasive species *G. costulatus*.

## Acknowledgments

We thank all the staff who participated in carrying out this study. We also thank EDCTP_CANTAM which funded this study and the CIRMF. Our thanks go to all the staff of RITMES at IHU MI in Marseille.

## Ethical considerations

This study has been approved by Gabon’s National Ethics Committee (CNER) under number: PROT 0014/2023/PR/SG/CNER.

## Funding

This work was supported by “EDCTP_CANTAM3”, “CIRMF” and a grant from the French Government managed by the National Research Agency under the “Investissements d’avenir (Investments for the Future)” programme with the reference ANR-10-IAHU-03 (Méditerranée Infection), by the Contrat Plan Etat-Région and the European funding FEDER IHUPERF.

## Supporting information Captions

**Table.S1.** Primers used to identify snails and detect parasites.

**Table S2.** Representative table of all the sites surveyed in Haut-Ogooué and Ogooué-Lolo.

**Figure S4.** Phylogenetic tree showing the position of the *Schistosoma haematobium* sequence and distribution of infected snail species. A- Tree constructed using COI gene sequences. B- Tree built with ITS2 gene sequences. The tree was constructed using the maximum likelihood method based on the Kimura. C- Geographical distribution of the snail species positive for *Schistosoma haematobium* by PCR.

**Figure.S5.** *Bulinus truncatus* and *Bulinus forskalii* distribution in Gabon. (A)- PCA of the LogScore values (LSV) of the *Bulinus truncatus* by sites (green=Mongouet, red=Libongui); (B)- Graphical representation (PCA), determining the classification of the LSV of the *Bulinus forskalii* by localities (blue= Okondja; green=Lastoursville; red=Ayanabo).

## Notes

### Competing Interest Statement

The authors have declared no competing interest.

